# Security and Privacy Concerns Regarding Genetic Data in Mobile Health Record Systems: An Empirical Study from Turkey

**DOI:** 10.1101/678912

**Authors:** Özlem Özkan, Yeşim Aydin Son, Arsev Umur Aydinoğlu

## Abstract

With the increasing use of genetic testing and applications of bioinformatics in healthcare, genetic and genomic data needs to be integrated into electronic health systems. We administered a descriptive survey to 174 participants to elicit their views on the privacy and security of mobile health record systems and inclusion of their genetic data in these systems. A survey was implemented online and on site in two genetic diagnostic centres. Nearly half of the participants or their close family members had undergone genetic testing. Doctors constituted the only profession group that people trusted for the privacy of their health and genetic data; however, people chose to limit even their doctor’s access to their genetic/health records. The majority of the respondents preferred to keep full access for themselves. Several participants had negative experience or preconceptions about electronic health records: the medical reports of 9.7% of the respondents had been used or released without their consent, 15.1% stated that they avoided being tested due to violation risks, and 3.5% asked their doctors to enter a less embarrassing health status in their records. The participants wanted to see some regulations and security measurements before using any system for their health/genetic data. In addition, significantly more participants stating that storing genetic data in a mobile system was riskier compared to other health data. Furthermore, the comparative analysis revealed that being young, being a woman and having higher education were associated with having greater privacy concerns.

## Introduction

In recent years, we have witnessed an impressive increase in DNA sequencing as it became much more affordable. It is projected that many citizens in developed countries will have their genomes sequenced within the next ten years (Ayday, De Cristofaro, Hubaux, & Tsudik, 2015); therefore, it is not hard to predict that the amount of genetic information included in health records will continuously increase. The inclusion of genetic/genomic information in electronic records can have a great impact on personalised healthcare by informing the physicians and patients about disease risks, differential diagnosis, right doses of drugs, as well as assisting in the selection of an effective treatment and preventive actions (McGuire et al., 2008). Even today, genetic testing is utilised more often for the diagnosis of increasing number of diseases; thus, integration of genetic test results or personal genomic data into electronic health records (EHRs) is an emerging issue. However, there are still many unanswered questions about where, when, and how to manage genetic/genomic data in EHRs (Shoenbill, Fost, Tachinardi, & Mendonca, 2014). Current EHR systems handle genetic data as any other laboratory tests; however, EHR systems should be redesigned to be more efficient and secure for genetic/genomic data (Shoenbill et al., 2014); since the privacy and confidentiality of genetic/genomic data presents unique challenges compared to other personal data due to its specific features, such as uniqueness, predictive capability, impact on other family members (Alahmad, Hifnawy, Abbasi, & Dierickx, 2016), temporality, ubiquity, and ease of procurement (Ayday, Raisaro, Mclaren, Fellay, & Hubaux, 2013; McGuire et al., 2008).

There are many ethical debates on the privacy and confidentiality of genetic information and potential to discriminate based on genetic background, such as employing or insuring individuals (Hoffman, 2017; Joly, Ngueng Feze, & Simard, 2013; Knoppers & Godard, 1998; Mohammed et al., 2017; Sherwin & Simpson, 1999; Sommerville & English, 1999). However, there are no comprehensive security solutions for protecting the privacy of genomic data. Even though several de-identification and aggregation techniques have been offered for the protection of privacy and security of EHR systems, their use in personal genomic data is limited since genome itself is an ultimate identifier for an individual (Malin, 2005). Today, privacy of genomic data is still an issue, but the literature contains a limited number of studies that propose solutions (Akgün, Bayrak, Ozer, & Sağıroğlu, 2015) despite the considerable amount of research reporting individuals’ concerns regarding the privacy and confidentiality of their genetic data (Alahmad et al., 2016; Heath, Ardestani, & Nemati, 2016; Henneman et al., 2013; Hietala et al., 1995; Jamal et al., 2014; Oliver et al., 2012; Sanderson et al., 2016). According to the general conclusions of these studies, people are mostly aware of the benefits of genome sciences, but they are concerned about the privacy of their data.

Mobile health applications are the fastest-growing and most popular tools among healthcare technologies. Currently, there are 325,000 mobile health applications in major application stores (Pohl, 2017). Although there are still privacy and security concerns regarding these applications (Arora, Yttri, & Nilse, 2014; Atienza et al., 2015; Luxton, Kayl, & Mishkind, 2012; Martínez-Pérez, de la Torre-Díez, & López-Coronado, 2015), it is projected that the global revenue of mobile health devices and services will be around 35.8 billion dollars by 2020 (Statista, 2017). Problems regarding the security and privacy of health data on mobile platforms should be addressed and concerns should be reduced before a system can be widely adopted. However, the design of mobile health applications still lacks features that would overcome the concerns of users, and applications with low security measures continue to be released to the market (Martínez-Pérez et al., 2015).

The international literature on people’s views and attitudes concerning genetic confidentiality is very extensive, but such studies are lacking in Turkey. Therefore, in this research, we aimed to develop and conduct a survey in Turkey to reveal the conditions that have the potential to convince people to record their genetic data in mobile applications. To identify the critical design elements of applications, we inquired about the preferences of users regarding what they would share and what kind of security protection they would prefer to ensure the security of their genetic data in these applications. Online banking safeguards were proposed as the gold standard for mobile information management, and it was investigated whether these security measures were also trustworthy for the management of health and genetic data in mobile applications. Finally, we elicited participants’ views concerning whether there was any difference between the privacy of personal, genetic and health data.

## Methods

### Pilot Study for the Assessment of the Survey

The survey was conducted in Turkish as it is the participants’ native language. A pilot study was performed with 20 participants to assess both timing and appropriateness of the survey items for the Turkish participants. After the pilot study, we conducted a follow-up interview with the participants to inquire about the clarity of the questions and based on their responses, we revised three questions. We also removed one of the questions from the instrument since the results showed that it was not informative. Hence, the final version of the survey consisted of 26 questions under three main categories: The first part covered demographic questions about city of residence, year of birth, gender, educational level, income, and computer and smartphone literacy levels. The second part aimed to obtain information about the participants’ level of awareness regarding security tools and their general online banking experience. The last part of the survey investigated the participants’ level of awareness, attitudes and experience concerning data security and privacy, and management of genetic and health data in mobile applications. Three questions were directly taken from two external resources (Canada Health Infoway & EKOS Research Associates, 2007; Princeton Survey Research Associates, 1999) and four questions were modified to address issues about genetic data.

### Data Collection

Data collection was undertaken using two methods: online and in-person. A snowball sampling method was used for online data collection between May 5, 2015 and August 2, 2015. An e-mail including the survey link was sent to university students and shared on Facebook pages with more than 15,000 users consisting of the students and staff of Middle East Technical University and people from the vicinity of the university. The participants were specifically asked to distribute the survey to people who or whose family member had undergone genetic testing. A total of 124 people responded to the surveys. After the elimination of 19 incomplete surveys, the remaining 105 were included in analysis. In order to reach more participants with different demographic profiles and increase the number of participants who were familiar with genetic testing, in-person data collection was implemented in two centres: a private genetic diagnostic centre and a medical genetic department of a university hospital. Sixty-nine people responded to the on-site surveys; thus, the total number of completed surveys was 174.

### Data Analysis

We used SPSS (version 23.0.0) for statistical analysis. Descriptive statistics were reported as frequencies. Pearson’s chi-square test was used to test whether there were any differences within the following seven groups: gender, age (≤ 35 versus > 35 years), educational level (university degree versus other), income [≤ 2000 Turkish Liras versus > 2000], computer literacy (< 4 versus ≥ 5), smartphone literacy (< 4 versus ≥ 5), and genetic testing [tested (himself/herself or a family member versus non-tested]. Post-hoc achieved powers were computed using G-Power 3.1.9.2. (Erdfelder, Faul, & Buchner, 1996). Cronbach’s alpha coefficient was used for reliability analysis (Pallant, 2013).

## Results

The internal consistency of the instrument was tested using Cronbach’s alpha coefficient for the analysis of reliability. Ideally, Cronbach’s alpha coefficient is expected to be greater than 0.7 (Pallant, 2013), and the current scale was found to meet the required value (0.72).

### Demographics of Respondents

The sample consisted of 174 people (100 women, 74 men) from 21 different cities in the Republic of Turkey. The average age of the participants was 34.09 (± 8.98). The average monthly income of the participants was between 2,001 and 4,000 TL. In Turkey, the hunger limit was 1,257 TL and the poverty limit for a family of four was 4,094 TL in 2015 (“Ocak 2015 Açlık ve Yoksulluk Sınırı,” 2015). The participants were from various educational backgrounds, with the highest frequency belonging to bachelor’s degree holders (50.6%), followed by graduate degree holders (35.1%). The percentages of other educational levels were as follows: high school graduates 10.3%, high school graduates 2.9%, and primary school graduates 1.1%. Most of the participants stated that they had an above-average (70.7%) or average level of computer literacy (20.7%), and similar rates for smartphone literacies: 73.6% for above-average level and 13.8% for average level.

### Level of Knowledge and Experiences on Health and Genetic Data

In the online survey, we reached 16 people who or whose family member had previous experience with genetic testing. In addition, 69 people from the two genetic testing centres completed the questionnaires. Thus, in the sample pool, the total number of people who or one of whose family member had previous personal experience with genetic testing was 85 (48.9%). Furthermore, to acquire the views of other participants who had not taken a genetic test before, the following question was added to the survey: “What would you do if you were offered genetic testing?” As a result, 96.6% of the non-tested participants reported that they would take a test if necessary.

A great majority of the participants (60.9%) stated that they did not know who had the right to access their medical records. Only a small number of the participants (7.4%) believed to have comprehensive knowledge on the topic. When asked about their knowledge on genetic science, 47.6% of the respondents indicated that they knew nothing or had very little knowledge. The rest of the participants (52.4%) had either average or above-average level of knowledge in this area.

Three items in the questionnaire (Q17-Q19) aimed to reflect participants’ experience about the sharing of their health data. Seventeen participants (9.7%) responded that their medical records had previously been inappropriately used or released without their consent (See Table 1).

**Table 1.** Responses to the item, “Have your medical records ever been inappropriately used or released without your consent?”

|  | n | % |
| --- | --- | --- |
| Yes, my data has been shared with a third party without my consent. | 4 | 2.3 |
| Yes, my data has been shared with my employer/my insurance company without my consent. | 11 | 6.3 |
| Yes, my data has been used in research without my consent. | 2 | 1.1 |
| No / I don’t know. | 156 | 89.7 |
| No response | 1 | 0.6 |

In addition to the breach of medical confidentiality, 15.1% of the respondents stated that they had avoided being tested to prevent others from accessing their results. Moreover, six participants (3.5%) asked their doctors not to write their symptoms/diagnosis in their medical records or enter a less embarrassing alternative rather than the actual condition (Table 2).

**Table 2.** Responses to the item, “Have you ever asked a doctor not to write your health problem in your medical records or to provide a less embarrassing diagnosis?”

|  | n | % |
| --- | --- | --- |
| Yes, I have asked a doctor not to include my health problem in my records. | 1 | 0.6 |
| Yes, I have asked a doctor to provide a less embarrassing condition for my records. | 5 | 2.9 |
| No, I haven’t. | 166 | 95.4 |
| No response | 2 | 1.1 |

## Attitudes Towards Health and Genetic Data Sharing

We observed that the participants were sensitive about sharing their health/genetic data with third parties and they thought some regulations were needed for the protection of their privacy. Question 21 (Q21) concerned the level of access rights regarding genetic data included in medical records, and the majority of the participants (94%) responded to this question by stating that they should have full access. Approximately half of the participants stated that their children (57.5%), parents (55.7%), doctors (52.5%), spouses (50.3%), and other doctors or hospital staff (45.9%) should have limited access rights. Lastly, a considerable number of participants did not want to give any access rights to their neighbours/friends (83.9%), drug companies (83.2%), employers (81.4%), close relatives (65.4%), insurance companies (62.2%), or pharmacies (59.4%).

The responses to the item (Q24), “Do you trust the following stakeholders to keep your genetic and medical data private?” revealed that for the majority of the participants (61.2%), doctors were the only trustworthy providers. The least trusted were insurance companies (82.6%), followed by information technology specialists (71.3%), the government (68.6%), pharmacists (63.9%), and nurses and other hospital staff (53.8%) (Table 3).

**Table 3.**
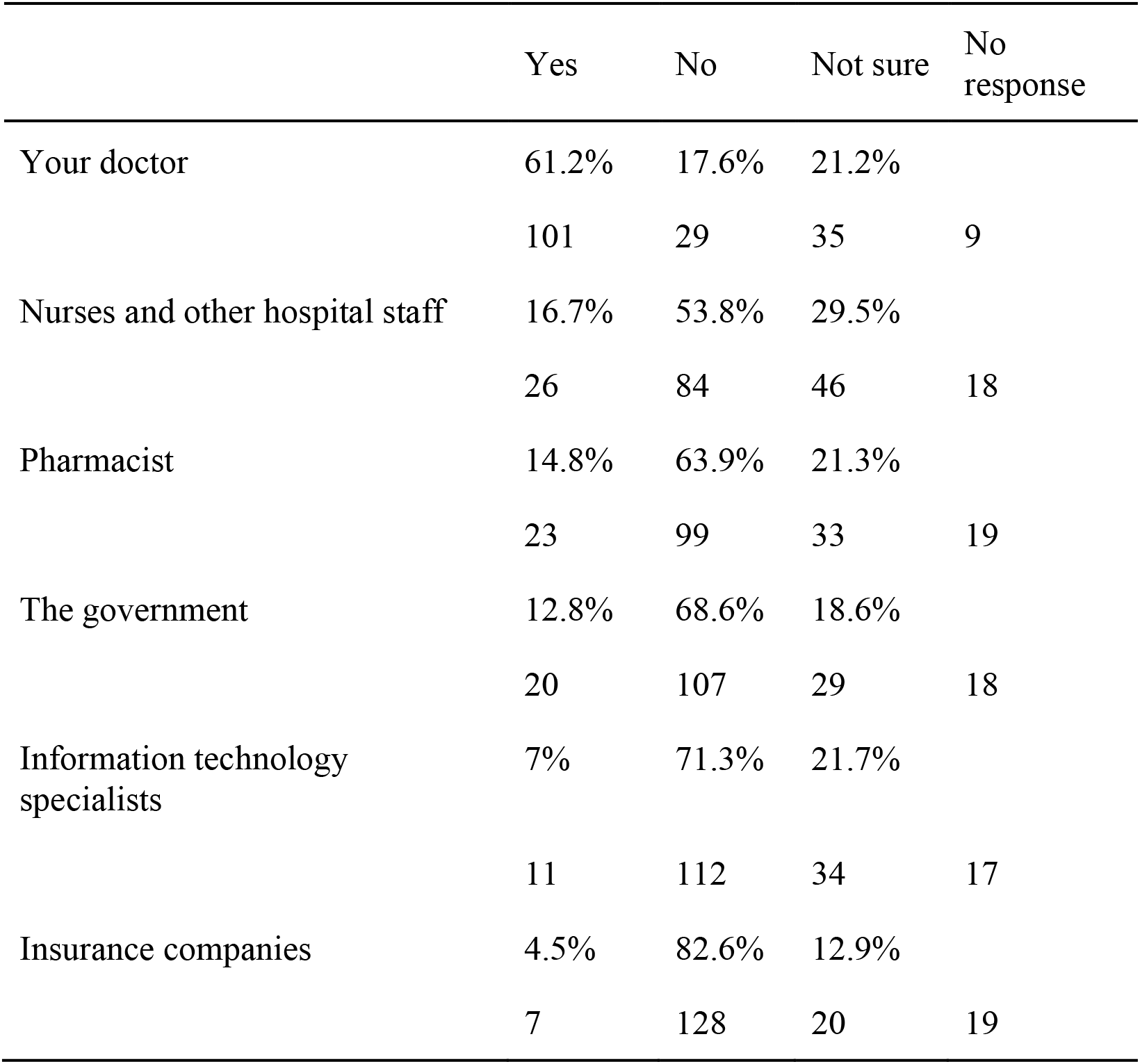
Responses to the item, “Do you trust the following stakeholders to keep your genetic and medical data private?”

Four regulations were proposed to the participants for the protection of the confidentiality and privacy of genetic data included in their electronic records (Q22). A five-point Likert-scale was used for the evaluation of this question, and according to the results, three of the four suggested regulations were found to be potentially effective (Options 1, 3 and 4). Option 2 was neither supported nor rejected. Details are given in Table 4.

**Table 4.** Respondents’ views about the effectiveness of regulations proposed to protect their privacy and confidentiality

| Options |  | Sum of 1-2 rates | Sum of 3-5 rates | Not sure | No response |
| --- | --- | --- | --- | --- | --- |
| C3: Using trustworthy security systems that use passwords and encrypted data on the device where your information is stored | %<br>n | 10.8%<br>17 | 82.8%<br>130 | 6.4%<br>10 | 17 |
| C4: Having the option to see when and by whom your records are retrieved | %<br>n | 14.5%<br>23 | 79.8%<br>127 | 5.7%<br>9 | 15 |
| C1: Ensuring that doctors and healthcare providers who need access to your medical information only use data that does not contain any personal identity information | % | 22.3 | 65.6% | 12.1% |  |
|  | n | 35 | 103 | 19 | 17 |
| C2: Having the option to see, correct, or even delete your medical records | % | 35.9 | 49.3% | 13.2% |  |
|  | n | 56 | 77 | 23 | 18 |

### Views on Mobile Applications for Health/Genetic Data Management

We collected the participants’ views concerning the use of mobile applications for health/genetic data management. Q23 inquired the kind of information the participants would like to keep in a health record application installed on their smartphone. We also wanted to determine whether the participants were willing to keep their genetic data in their mobile health record application; therefore, we added “diseases in your family” to the options. The participants were allowed to choose more than one option for this question. All the options were chosen by more than 50% of the participants. The top six responses were allergies (84.2%), medication (83%), in case of emergency (ICE) number (81.9%), diseases and health problems (77.8%), operations (72.5%), and vaccines (71.3%). The option of diseases in your family had a lower response rate compared to the others, but it was still chosen by more than half of the participants (56%). The details of the responses are presented in Figure 1.

**Figure 1.**
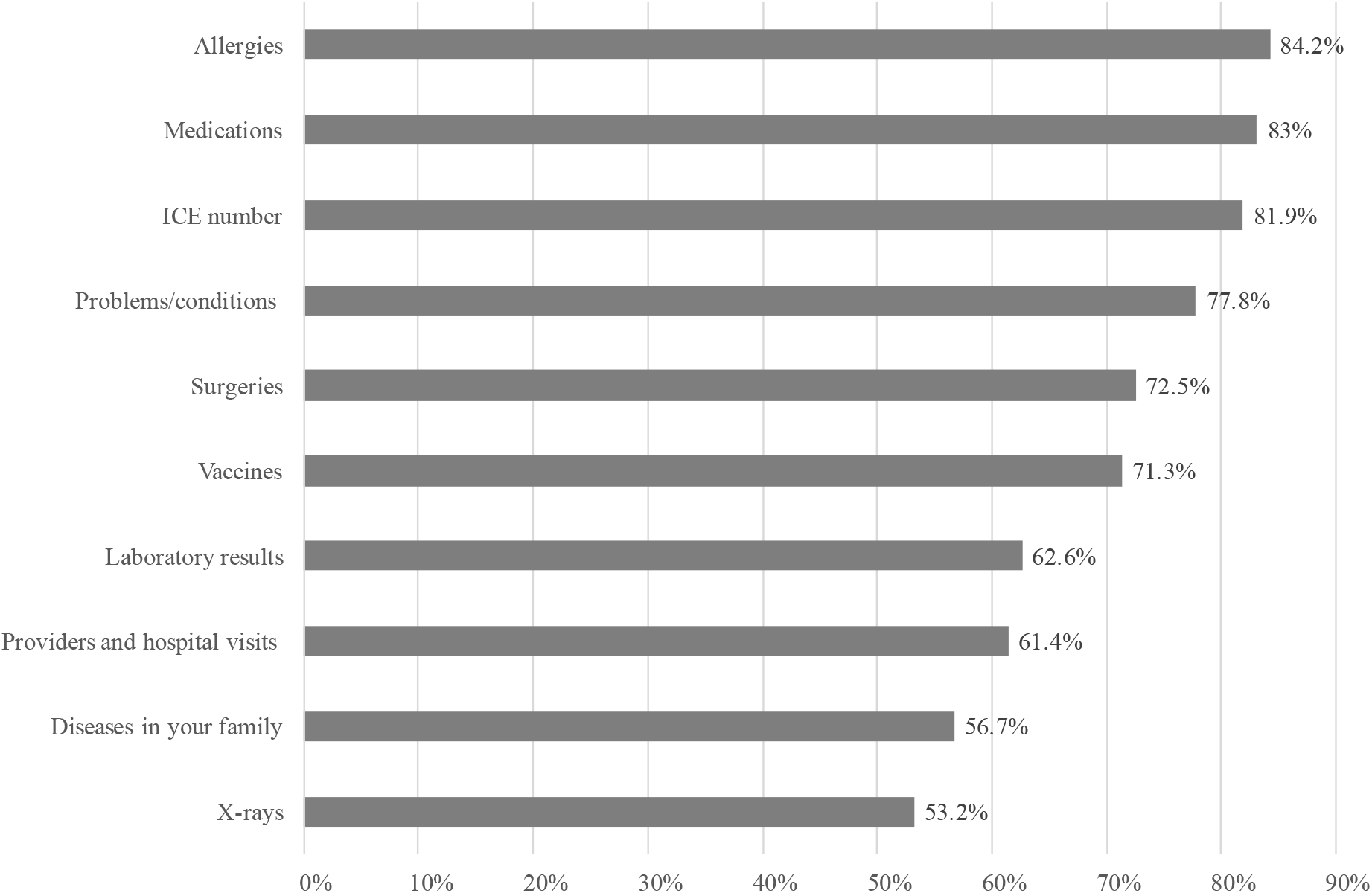
Types of information the users wanted to see in a mobile health record application

**Figure 2.**
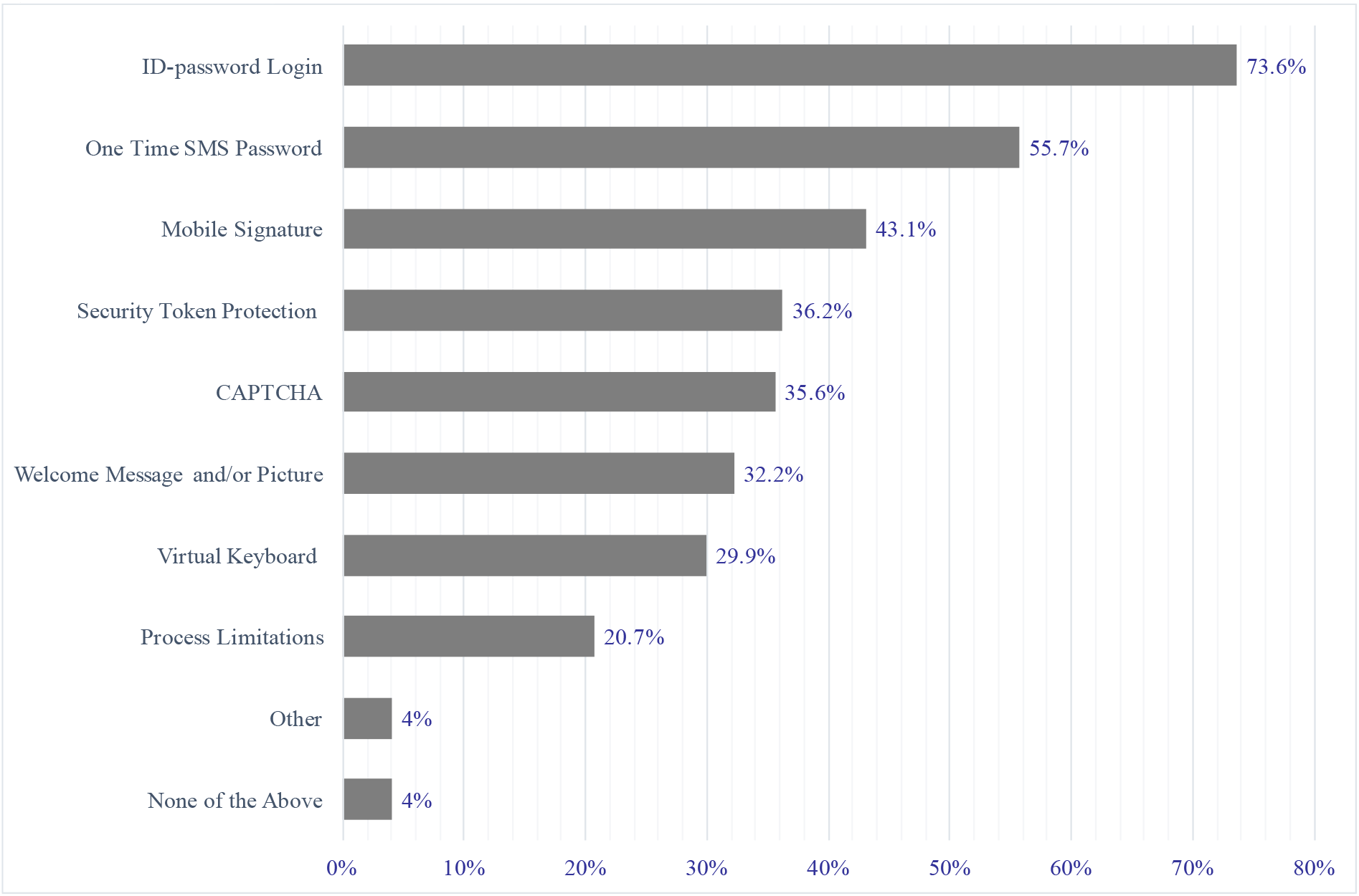
Security features the users would like to see in an internet-based health and genetic data record system

Q25 was related to the participants’ attitude concerning the privacy requirements of different types of information and risks involved in storing it in a mobile application. The responses to this question are presented in Table 5. **Error! Reference source not found.** The results demonstrated different attitudes towards health and genetic information. A significantly (*P* = .00) higher number of participants considered that genetic information was at a higher security risk (62.2%) than health information (44.6%) stored in mobile applications, and the respondents were concerned about the security of their genetic information nearly as much as their identity and personal information.

In terms of online banking, most participants (87.4%) reported that they had used these services before, and only 16.1% of these participants considered online banking safeguards to be insufficient for protecting the security of their information. Moreover, eight participants (4.8%) previously had negative experience when using online banking, one of whom still believed that using online banking was secure and three were not sure about it. The responses to Q25 showed that the majority of the participants either thought that none of their information could be safely stored on mobile platforms or were not sure about the risks involved. Bank account information was at the top of the list, being chosen by 81.5% of the participants. Despite these negative views, our analysis showed that almost all the participants had used or were using online banking systems (Table 6).

**Table 5.**
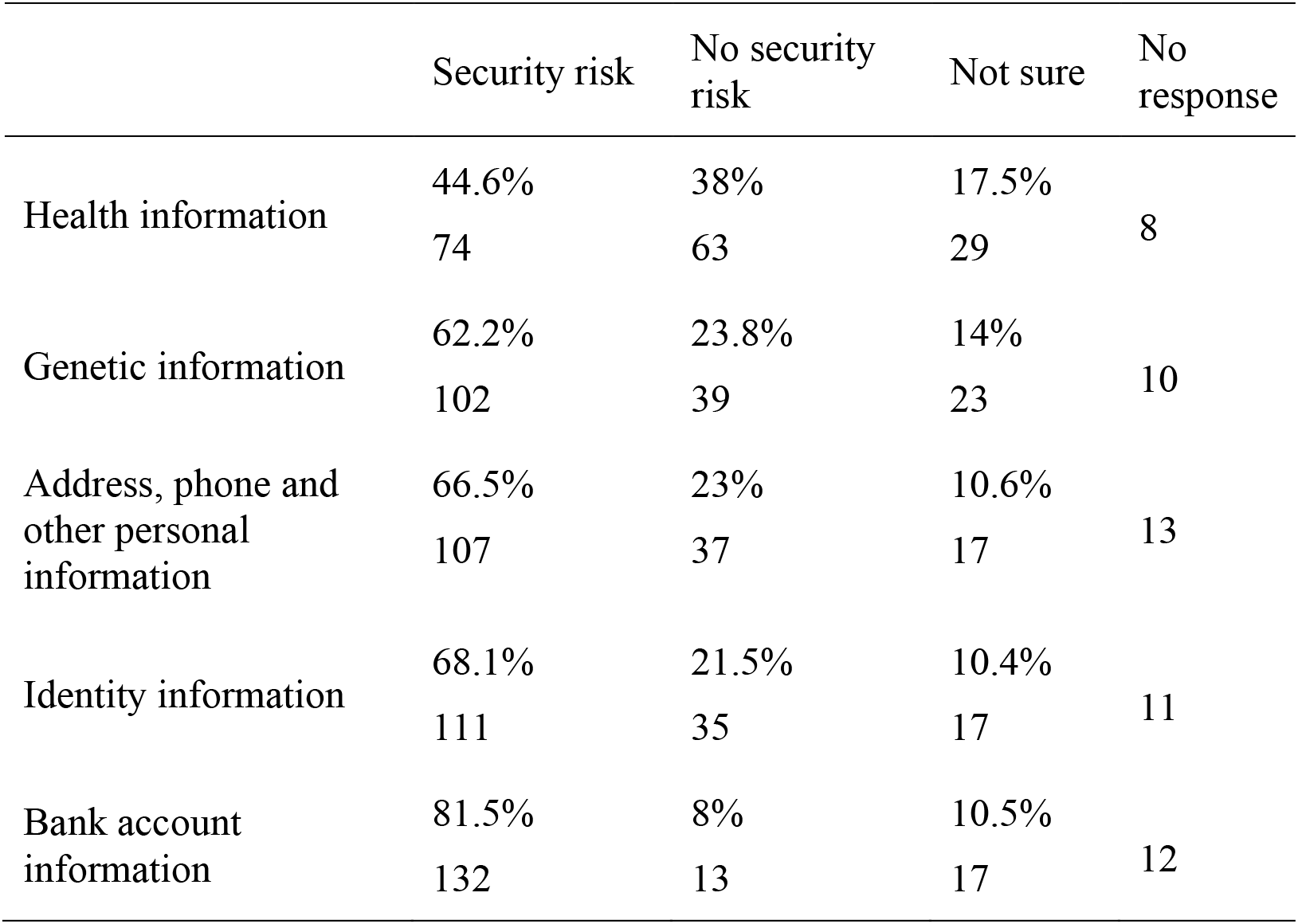
Responses to the item, “What do you think about the security risks of storing the following information in a mobile application?”

|  | Security risk | No security risk | Not sure | No response |
| --- | --- | --- | --- | --- |
| Health information | 44.6%<br>74 | 38%<br>63 | 17.5%<br>29 | 8 |
| Genetic information | 62.2%<br>102 | 23.8%<br>39 | 14%<br>23 | 10 |
| Address, phone and other personal information | 66.5%<br>107 | 23%<br>37 | 10.6%<br>17 | 13 |
| Identity information | 68.1%<br>111 | 21.5%<br>35 | 10.4%<br>17 | 11 |
| Bank account information | 81.5%<br>132 | 8%<br>13 | 10.5%<br>17 | 12 |

**Table 6.** Cross tabulation of the participants’ views on storing bank account information in mobile applications and their experience with online banking

|  | Bank account information |  |  |  |
| --- | --- | --- | --- | --- |
|  | Security risk | No security risk | Not sure | Total |
| Have you ever used online banking? |  |  |  |  |
| Yes | 114 | 12 | 15 | 141 |
| No | 18 | 1 | 2 | 21 |
| Total | 132 | 13 | 17 | 162 |

Q20 was directed to the participants to determine their preferences related to the security measures in an application that would store their medical and genetic data. The participants were allowed to choose more than one option for this question. Almost all the respondents (96%) preferred the application to have at least one security feature, with the prominent responses being ID password login (73.6%), one-time password (OTP) over SMS (55.7%), and mobile signature (43.1%) (Figure 2). In addition to the options we provided for online banking security, we included the ‘other’ option for participants to make their own suggestions. Three participants suggested using fingerprint and one participant suggested voice authorisation system as a security feature.

**Figure 2.**
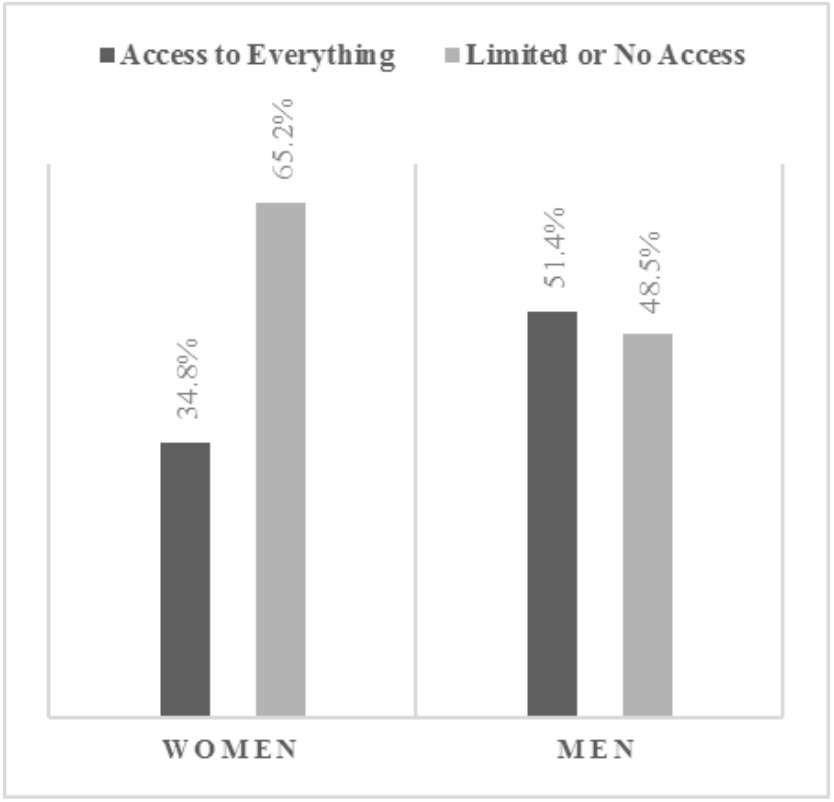
Comparison of the views of women and men regarding the rights of their spouse to access genetic data in their medical records (The post-hoc computed achieved power for w=0.3, alpha=0.05 and n=165 was 97.0%)

### Differences Between Groups

Gender, age, educational level, and computer and smartphone literacy levels were found to have a significant effect on the views of the participants. No significant differences were observed in the income and experience of the genetic testing groups. Gender was found to significantly affect the participants’ views concerning the access rights of their spouses (*P* = .04). Unlike men, women tended to give limited or no access rights to their spouse (Figure 3).

A significant difference (*P* = .02) was found between university graduates and those from other educational backgrounds in terms their views on the rights of their doctor to access their medical and genomic data. Unlike the participants with a lower level of education, most participants with university or above degrees preferred their doctors to have limited or no access to their medical and genomic data (Figure 4).

**Figure 4.**
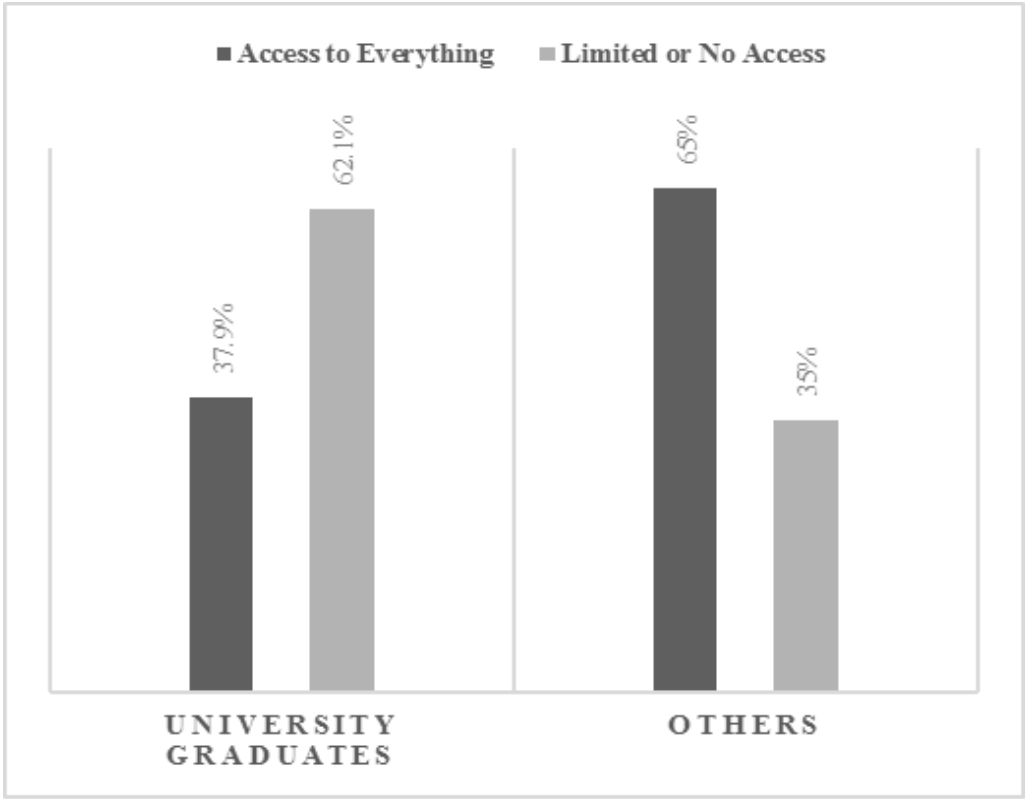
Comparison of the views of university graduates and those from other educational backgrounds regarding the rights of their doctor to access genetic data in their medical records (The post-hoc computed achieved power for w=0.3, alpha=0.5, and n=160 was 96.6%)

When the chi-squared test was repeated for the age groups, a significant difference was seen in the preferences of access by third parties for under the age of 35 (*P* = .00). Although the majority of the age groups tended to give their children limited or no access to their medical and genomic data, participants younger than 35 had more concerns about sharing their data with their children (See Table 7).

**Table 7.** Participants’ views on their children’s access to their genetic and medical data by age group (The post-hoc computed achieved power for w=0.3, alpha=0.01 and n=154 was 87.4%).

|  | Access to everything | Limited or no access | Not sure | No response | P |
| --- | --- | --- | --- | --- | --- |
| ≤35 | 24 | 99 | 4 | 16 | .00 |
| >35 | 15 | 16 | 0 |  |  |

Furthermore, the intra-group comparison of computer and smartphone literacy groups demonstrated a significant difference of opinion between the participants regarding their doctors’ right of access to their genetic and medical data. People with a high level of literacy in both areas chose to give their doctors limited or no access compared to those with a lower level of literacy (Table 8).

**Table 8.** Participants’ views on their doctor’s access to their genetic and medical data by level of computer and smartphone literacy. (The post-hoc computed achieved power for w=0.3, alpha=0.01, and n=160 was 88%)

|  | Access to everything | Limited or no access | Not sure | No response | P |
| --- | --- | --- | --- | --- | --- |
| Low Computer Literacy | 27 | 19 | 1 | 12 | .00 |
| High Computer Literacy | 39 | 75 | 1 |  |  |
| Low Smartphone Literacy | 23 | 17 | 1 | 12 | .02 |
| High Smartphone Literacy | 43 | 77 | 1 |  |  |

## Discussion and Conclusion

Our results showed the respondents’ sensitivity about sharing health/genetic data with third parties. They wanted to see regulations and security measures for the protection of their data. The participants chose to trust only their doctors for the privacy of their health and genetic data, but they chose to limit even their doctors’ access to their genetic/health records. Although most participants did not oppose keeping their health and genetic data in a mobile health/genetic data management application, significantly more participants thought that their genetic data was at a higher security risk in such applications. We also found that privacy concerns were greater among the young, female, and more educated respondents.

We also asked the participants some questions to understand their concerns about the current health record system. Three questions (Q17-Q19) related to the participants’ experience had been used in a previous survey conducted in 2011 in Turkey (Özkan, 2011). Although, that survey had a higher number of participants (596), the participant profile was very similar in terms of age (average: 28.6) and educational level (university or above: 87.3%). There was a noticeable difference between the two surveys in the percentages of ‘yes’ responses to the three questions listed (the results of the current survey are given in parenthesis): 0.8% (9.7%) reported that either their or one of their family member’s medical records had been inappropriately used or released without their consent, 12.5% (15.1%) avoided being tested in case someone might see their results, and 1.5% (3.5%) asked their doctors to include a less embarrassing alternative in their medical records rather than their actual condition. We find these results alarming since they support the idea that patients are inclined to postpone or give up treatment or change the circumstances in which their illness occurred or withhold certain details because they have concerns about confidentiality. An urgent action plan is needed to establish greater public confidence about the confidentiality and privacy of health, genetic and other records.

Another important finding of the survey was the observation that only a small number of participants had comprehensive knowledge regarding the rights of access to their health records. Unfortunately, the remaining participants either did not know anything or have doubts about the issue. When asked about their preferences about the access rights, most believed that they should be the only people with full access to their medical and genetic data. These results indicate that people would like to have sole control over using their data and favour a self-controlled rather than a centralised, multi-user system. An exploratory survey conducted in Saudi Arabia (Alahmad et al., 2016) contained similar questions about access rights. Although there was no limited access option in the Saudi questionnaire, the results were parallel to ours. The Saudi participants preferred to give their doctors and themselves access to their medical data but refused to do the same for insurance companies. The health systems, policies, and regulatory frameworks of both countries should be discussed to see the reasons behind the similarity of the results obtained from these two countries.

Most of the participants in this study (79.8%) also wanted to have the option to see when and by whom their records were retrieved. Similarly, Atienza et al. (2015)(Atienza et al., 2015) reported that the participants’ concerns about privacy and security of mobile health data were connected with when and by whom information was accessed and seen. Our overall results suggest that the system should allow tracking access and ask for the patient’s permission prior to releasing or distributing their medical data or sharing anonymous information that does not contain any personal details. Furthermore, an effective health information system should allow the user to hide sensitive information from users that are not authorised by the data owner.

According to our results, significantly more men preferred to give their wives full access to their genetic and medical data. This may have several reasons. In the literature, studies have shown that women are less willing to participate in genetic studies or allow storage of their genetic data than men (Espeland et al., 2006; Matsui, Kita, & Ueshima, 2005). One reason for this may be the privacy concerns of women about genetic tests. However, the results of similar studies on gender differences are not consistent (Khan A, Capps BJ, Sum MY, Kuswanto CN, & Sim K, 2014). For example, Mezuk, Eaton, and Zandi (Mezuk, Eaton, & Zandi, 2008) found no association between genders about consent to donate a biological sample or allow genetic testing or storage of that sample while Green et al. (Green et al., 2006) reported that it was men that mostly denied private companies access to their DNA.

Another reason behind women being less willing to share their medical data could be that they have more information in their medical records than men since they are admitted to healthcare facilities more often (Ashley, 2010; Brett & Burt, 2001; Goldstein, 2010). Excluding prenatal examinations, statically more women visit hospitals for psychiatric and chronic diseases (Ashley, 2010; Bates, 2011; Goldstein, 2010). So, one justification to our observation can be men having a lower level of concern about their health or health records since they experience fewer medical conditions. Lastly, there can be cultural or country-specific reasons for the differences of views between genders. In many countries, women still cannot express themselves freely since they are under the pressure of men and society in the majority of areas from business and economy to family (Kandiyoti, 1977, 1988). The discrimination between men and women may result in the latter feeling less comfortable sharing their private information with their spouses.

Parallel with our results, in the literature, younger age has been associated with greater privacy concerns (Khan A et al., 2014). Oliver et al. (Oliver et al., 2012) provided a very interesting interpretation for the difference between the age groups. According to the authors (Oliver et al., 2012), the reason for older people having less concerns of old people on regarding the commercialisation of their DNA is their belief that it would take years before a person can be identified from their DNA on the Internet, and this would not probably be possible in their lifetime. Despite being an unusual explanation, this may also be the underlying reason for our results. The situation was similar for educational level. McGuire et al. (McGuire et al., 2011) reported that university degree holders were more likely to choose restricted data sharing similar to our participants’ tendency to limit their doctors’ access to their data. Our study also showed that different groups had varying perceptions and views which should be taken into consideration when designing genetic-health data systems. As also suggested by Aro et al. (Aro et al., 1997), “age, education and gender related differences in acceptance of genetic testing which need to be taken into account when considering screening programs and informing the public”.

Our findings showed that the level of trust in terms of ensuring the confidentiality of genetic and medical data ranged from higher to lower as follows; doctors, pharmacists, nurses and other hospital staff, the government, information technology specialists, and lastly insurance companies. In general, the participants tended to trust people who were directly involved in their treatment more than the government, insurance companies, or information technology staff. Moreover, the results showed that people trusted only their doctor about their health and genetic data, which is promising as no quality healthcare service can be provided when there is no trust in the provider. The analysis of the same question showed that information technology specialists are one of the least trusted groups by the participants. However, health information technology professionals have a key role in protecting the security of genetic data and minimising breach risks when designing related systems (Shoenbill et al., 2014). Very recently, many significant steps have been taken on personal data privacy legislation in Turkey. The first Turkish Personal Data Protection Law (numbered 6693) was published on March 24, 2016 (“Kisisel verilerin korunmasi kanunu,” 2016), which brings about new sanctions and punishments for those who do not protect privacy of data. This may not necessarily reduce people’s concerns about this issue yet, but considering that the law is relatively new, the actual outcomes will only be observed over time.

Nearly half of the participants or at least one of their family members had been genetically tested before the study and the majority of the remaining participants stated that they would take a genetic test if necessary. This result is valuable since the questions about sharing, storing and protecting genetic information provided an insight into the participants’ views based on their actual experience in addition to a hypothetical scenario. The responses also revealed that the participants had a different attitude towards their genetic information compared to other medical information, with more people finding it risky to store their genetic information in mobile applications. As in other types of health-related data, the participants were also not very willing to keep information about family diseases in an application. In contrast, Alahmad et al. (2016) found no significant difference in giving access to medical or genetic records. Our participants compared the value of genetic information to their identity and personal information, such as address and phone numbers. Many people also stated that a mobile health record application should have similar security protections such as OTP and mobile signature to keep their health and genetic information secure. This indicates that a mobile health record application with sufficient security protections have great potential for adoption by mobile users. Otherwise, as discussed by Heath et al. (Heath et al., 2016), “privacy concerns will have a negative influence on behavioural intentions to share genetic information”.

In conclusion, the results of our study show that people would like to have a system that will give them full control over their health and genetic data, make them feel safe and secure about sharing, hiding or even deleting their information. The system should also be flexible in terms of being adaptable to user preferences. All of these requests are pointing to a personal, self-controlled health application. A well-designed, patient-oriented, and secure mobile personal health record application would have the potential to be widely adopted and efficiently used for health and genetic data management.

## Limitations and Future Work

The current study has two main limitations: Firstly, collecting data from genetically tested patients was challenging since most people in the diagnostic centres either refused to participate in the study or did not fully complete the questionnaire. Approximately 85% of the incomplete questionnaires belonged to respondents with a high school or a lower level of educational background. This situation was similar in the online version of the questionnaire, with only 5.7% of the responders having a lower educational background. Therefore, we were not able to analyse the overall difference between the results according to the educational levels of the participants. Secondly, the sample size was limited and was not generalizable to the whole Turkish population. For future research, the survey can be administered to a sample that would reflect the Turkish public opinion. Moreover, a health record application including mobile genetic data can be designed addressing the concerns revealed in the study, and a usability analysis can be implemented with various user groups to elicit and compare their views. The results of such research would make a positive contribution to the development of applications that are secure and trustworthy.

## Funding

The author(s) received no financial support for the research, authorship, and/or publication of this article.

